# Direct Measurement of Dissipation in a Single Protein using Small Amplitude Atomic Force Microscopy

**DOI:** 10.1101/275065

**Authors:** S. Rajput, S. Talele, V. Ahlawat, VJ. Ajith, A. Roychoudhury, S. Kamerkar, S. Patil

## Abstract

In Krammer’s theory, stiffness and dissipation coefficient of a protein determine the rate of their conformational change. Using atomic force microscope, it is possible to measure viscoelasticity of a single protein, wherein it’s dissipative and elastic nature is directly and independently measured. Such measurements are performed, either by measuring the thermal fluctuations of the protein held under a constant force, or by providing small modulations to the protein by dithering the cantilever and measuring its response. In small amplitude approximation, where dither amplitude is comparable to persistence length of polymers, it is possible to measure the protein’s viscoelastic response accurately. We measured dissipation in I27 at extremely low pulling speeds (∼ 50 nm/s) and low dither frequencies (∼100 Hz). At these experimental parameters the dissipation is found to be ∼10^−5^ kg/s, well above the detection limit of conventional AFM and upper limit predicted by Benedetti et al. Our stiffness data clearly reveals unfolding intermediate of titin’s individual immunoglobulin units. The intermediate is elongation of folded domains by ∼8 Å, wherein two hydrogen bonds are broken between beta sheets. It was possible to measure this elongation in our experiments. The directly measured internal friction of unfolded polymer chain shows a scaling with tension on the chain. The measurements show that it is possible to measure internal friction in single molecules unambiguously using small amplitude AFM. It suggests that systematic experiments to unravel the relation between directly measured internal friction and folding rates of proteins are possible.

## INTRODUCTION

In last few decades, a number of experimental techniques have been developed to observe and manipulate matter at atomistic and molecular scales [1–10]. These developments have a lasting impact on the field of molecular biology as well as on molecular nanotechnology. Single macromolecules such as proteins, flexible polymers and polysaccharides are stretched and coil-to-stretched or folded-to-unfolded transitions under the application of external force are routinely observed. The central goal of these experiments is to mimic the molecular response under natural situations such as ligand-receptor binding, allosteric signalling and the stress induced conformational changes. They measure kinetics of unfolding of a protein and binding affinities of ligands to receptors.

Along with optical tweezers [3, 9–12], Atomic Force Microscopy has acquired a unique position in this quest due to its unprecedented spatial resolution in physiological conditions[5–7, 13–17]. In typical AFM experiment, mica or Au substrate is sparsely coated with the biological macromolecule and is placed in the liquid environment. A sharp probe attached to a cantilever spring is then brought close to a molecule. The protein is allowed to attach to it from either C or N terminus through nonspecific binding. The bending in the cantilever beam is measured as the protein is pulled away from the surface. The cantilever bending provides the amount of force applied on the protein as it is slowly stretched. The spectrum of force required to unfold protein at different pulling rates is recorded and used in either Monte Carlo simulations or theoretical calculations to draw a picture of energy landscape [18–22]. A large body of work exists in the literature that utilises this method to extract energy landscape of single macromolecules of biological interest including proteins and large polymers such as polysaccharide chains.

In aforementioned static experiments, wherein the cantilever is not oscillated while pulling at a protein, one is not able to utilise the phase sensitive detection methods which effectively separate signal from noise and enhance force sensitivity. These methods also allow measurements of shifts in cantilever’s frequency or amplitude along with phase information which can be quantitatively related to viscous and elastic components of the force [23–26]. This is rheology on single molecule scale. The viscous response is mainly due to the internal friction of protein while the elastic response is attributed to its stiffness.

The measurement of viscous and elastic components of protein’s response to external periodic perturbations is of relevance to all biochemical reactions involving conformational changes in protein. In Kramer’s rate theory, the ratio of dissipation coefficient *γ* of a protein to its stiffness *k* determines the time over which its shape is statistically uncorrelated[27]. This ratio 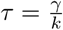, typically called relaxation time is a primary factor limiting the rates of biochemical reactions initiated through a application of mechanical strain on the protein. In this context, it is of paramount importance to determine the stiffness, and dissipation arising from internal friction directly and simultaneously. It can be measured through a small dithering of a single protein tethered to a cantilever. Its response, obtained using changes in cantilever’s amplitude and phase, can potentially provide such direct measurement of stiffness and dissipation.

While it is relatively straightforward to quantitatively estimate the stiffness of a single protein, the estimation of dissipation coefficient is marred with controversies. The primary source of this disagreement is due to the cantilever’s hydrodynamic drag in physiological buffers. The cantilever is approximated as a point mass in setting up its equation of motion. This approximation works well in ambient as well as ultra high vacuum conditions, but fails in situations where it is oscillated in liquid environments. Recently, Benedetti et al. have showed that consideration of geometric details of the cantilever is crucial for estimating the dissipation of single molecules [28]. Their improved analysis of the data suggested that molecular dissipation (∼10^−7^ kg/s) is much lower than the detection limit of AFM.

Here, we measured dissipation in a single polyprotien consisting of 8 IgG domains of muscle protein titin (I27) using AFM. We analysed our data using the method presented by Benedetti et al. We contradict their conclusion that the dissipation in single molecules is below the detection limits of AFM experiments. We demonstrate that this conclusion arises from inappropriate choice of experimental parameters. A carefully chosen operation regime which ensures true off-resonance conditions and extremely slow quasi-static pulling (∼ 50 nm/s) with better measurement bandwidth allows measurement of dissipation in single proteins. Further, both the dissipation and stiffness signals contain features from intermediate in I27. These are previously detected in static experiments and steered molecular dynamics simulations have suggested that it is a result of breaking two hydrogen bonds between *β* sheets and elongation of a folded domain by ∼8 angstroms. Our findings establish the ability of AFM to measure dissipation. We also show that the dissipation is strongly dependant on operation frequency and has surprisingly large value (∼10^−5^ kg/s) below 100 Hz and it drops to 10^−7^ kg/s at 10 KHz. In our measurements the intermediate encountered upon stretching is more clearly and repeatedly seen when the number of folded domains are more than four. The corresponding static signal, however fails to detect this intermediate reproducibly. The use of phase sensitive detection methods such as lock-in amplifiers thus allow measurement with better force sensitivity. The estimated smallest measurable viscous and elastic component are ∼ 10^−7^ kg/s and 0.8 mN/m respectively.

## MATERIALS AND METHODS

### Sample preparation and AFM measurements

Titin I27 polyprotein constructs containing eight identical domains in tandem were constructed from a plasmid similar to described in [29]. Expression and purification of the protein was done as described before [30]. PBS at pH 7.4 was used as the standard buffer for all experiments The protein sample (100 *µ*L) in PBS with a concentration of 10 mg/mL was adsorbed onto a freshly evaporated gold coated coverslip assembled in the fluid cell by incubating it on the substrate for 15 minutes at room temperature. The sample solution was washed three times with PBS to remove the excess un-adsorbed protein from the working solution. The unfolding experiments were performed using a commercial AFM (Model JPK Nanowizard II). Micro-fabricated commercially available gold coated cantilevers, with dimensions length = 300 ±5*µ*m, width = 35± 3*µ*m and thickness = 2 ± 0.5*µ*m, are used. The stiffness is calibrated using thermal noise method. The stiffness of the cantilever used in the measurement is 0.57 N/m. The range of resonance frequency is around 8.9 kHz in water. The cantilever is oscillated with free amplitude in the range of 0.5-2Å, at a frequency which is less than one third of resonance. The flatness of the frequency response in this region is ensured for off-resonance operation. The cantilever is held in place by a spring and its base is attached to a dither piezo. A sinusoidal drive voltage is supplied to dither using the lock-in SR830 (Stanford Research Systems). After completing the sample preparation protocol, using the cantilever deflection as signal for feedback control, the tip is auto-approached towards the surface. The point of approach was defined to be the tip applying below 100 pN of force on the sample during approach. Experiments were performed on ∼10 × 10 *µm*_2_ grid in a single run with total of 64 or 128 measurements. The attachment of protein terminus to the tip was achieved by non-specific interaction between the tip and sample. The tip was retracted from the surface at a constant speed of 40 nm/s. The stiffness and dissipation of the polyprotein construct I27 is measured while it is being unfolded.

**FIG. 1:**
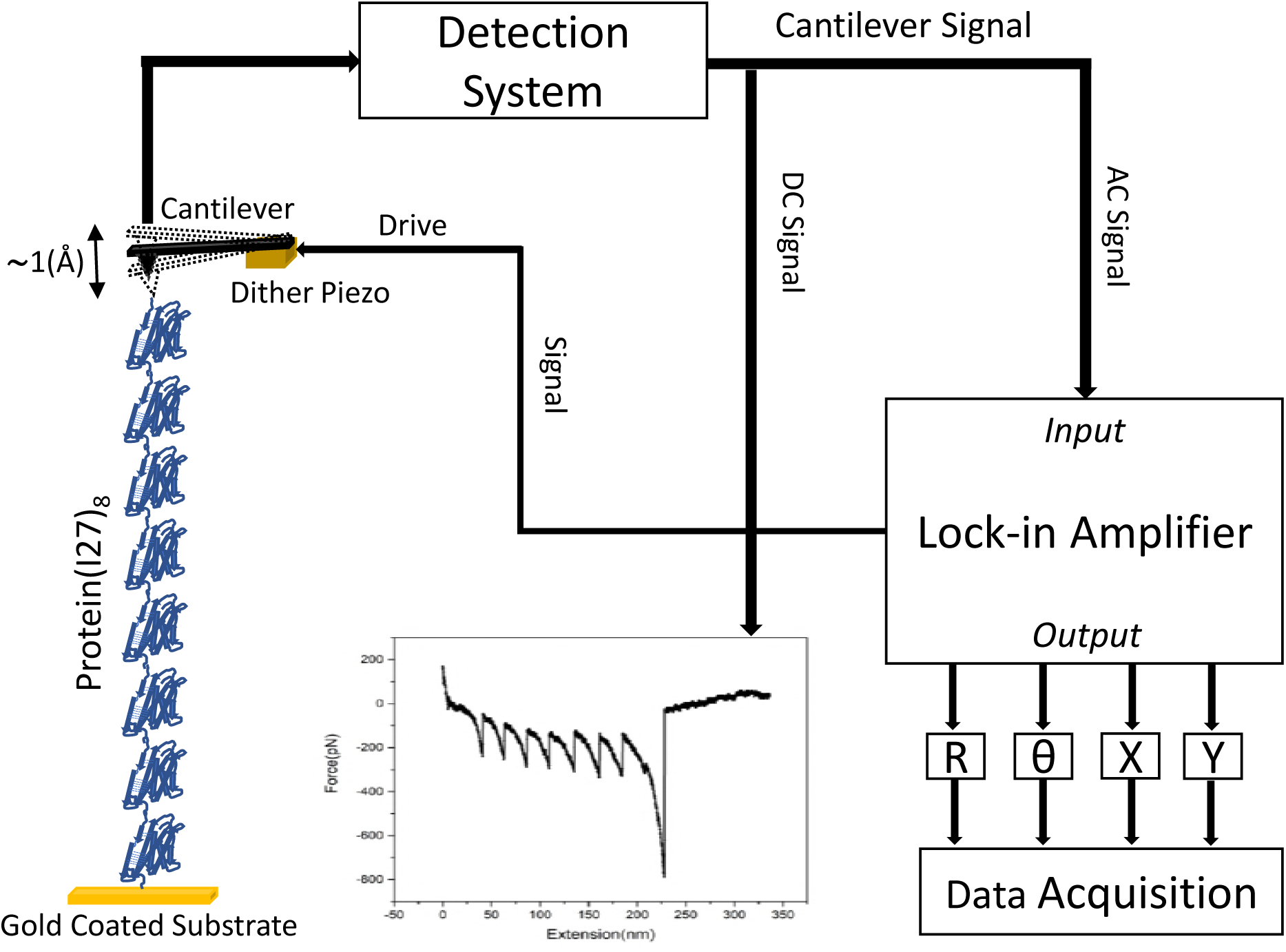
Schematic of the measurement. A small off-resonance dither (∼ 1 Å) is provided to the cantilever pulling on a protein. Along with a conventional saw-tooth pattern in force profile, we measure phase and amplitude of the tip oscillations using a lock-in amplifier. These signals are used for calculating dissipation and stiffness of a single protein in both folded and unfolded states.

### Data analysis methods

We discuss data analysis method employed in this work. While setting up the equation of motion, the cantilever assumed to be a point-mass attached to a massless spring of stiffness *k*_*c*_, and the viscous damping coefficient of *γ*_*c*_. This assumption may work in ultra high vacuum environments[42] and ambient conditions [43], but may lead to incorrect estimation of force in liquid environment[28].

In contrast to point-mass model approximation for cantilever dynamics, many theoretical treatments have considered geometric details of cantilever beam. This formalism is used for calibration of cantilever stiffness in liquid environments. The resulting equation of motion of the cantilever is fourth order partial differential equation in space and time [36–40]. To solve this equation, the boundary conditions at the free and fixed ends of the cantilever are used. Benedetti et al. have used this formalism in the context of quantifying the dissipation in single molecules measured by dithering the cantilever tip in off-resonance conditions. In this work, the data analysis used for calculating dissipation (*γ*_*i*_) and stiffness (*k*_*i*_) from measured quantities is based on this formalism. The equation of motion is

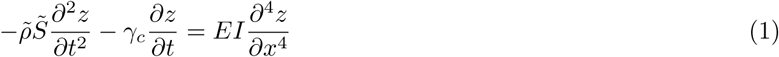

Where *z*(*x, t*) is cantilever displacement at position *x* from the base (*x* = 0) and at time *t*. The *z*(*x, t*) performs a simple harmonic motion and can be represented by a rotating vector having *R* having components *X* and *Y* given by *X* = *Rcos*(*θ*) and *Y* = *Rsin*(*θ*). The amplitude and phase of the oscillations is given by *R*_2_ = *X*_2_ + *Y* _2_ and *θ* = *tan*^−1^(*Y/X*). After applying suitable boundary conditions, the protein stiffness and dissipation can be calculated by recording X and of tip oscillations.

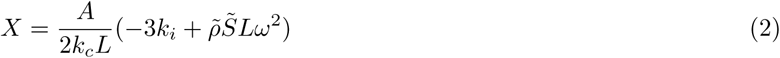

and

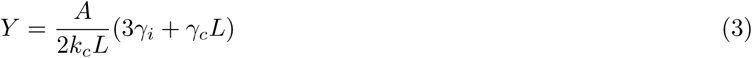

We record X and Y using lock-in amplifier and use equations 2 and 3 to estimate the stiffness (*k*_*i*_) and dissipation (*γ*_*i*_) of I27. 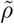 and 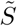 are modified density of silicon and cantilever cross section to account for effective viscous damping on the cantilever. *ω* and *A* are drive frequency and amplitude, respectively. *L* is cantilever length. *E* is elastic modulus and *I* is area moment of inertia determined by the geometry of beam.

## RESULTS

### Experimental Data

Figure 2 (a) and (b) show as received X and Y signals from lock-in amplifiers respectively. All four channels of data *X, Y, R* and *θ* can also be measured simultaneously. The measured amplitude and phase and those computed from *X* and *Y* are similar. The data shown here is representative of 100 similarly performed measurements. In experiments reported by Benedetti et al., the y-component of the amplitude (*Y*) is featureless while the domains are sequentially unfolded. The application of beam theory to estimate dissipation suggests that it depends exclusively on y-component of amplitude (Y). See equation 3. The featureless *Y* signal in their experiments leads them to conclude that dissipation in single molecules is below the detection limit of AFM. Note that in our measurements, we see clear modulations in the Y signal corresponding to unfolding events seen in the cantilever deflection data shown in Figure 2(c). The difference in experiments of Benedetti et al. and ours is the operation regime. We have used cantilevers with stiffness 0.6 N/m and resonance frequency of 8.9 kHz in water. Our drive frequency is 1.4 kHz, free amplitude is 1.3 Å and pulling speed is 40 nm/s.

**FIG. 2:**
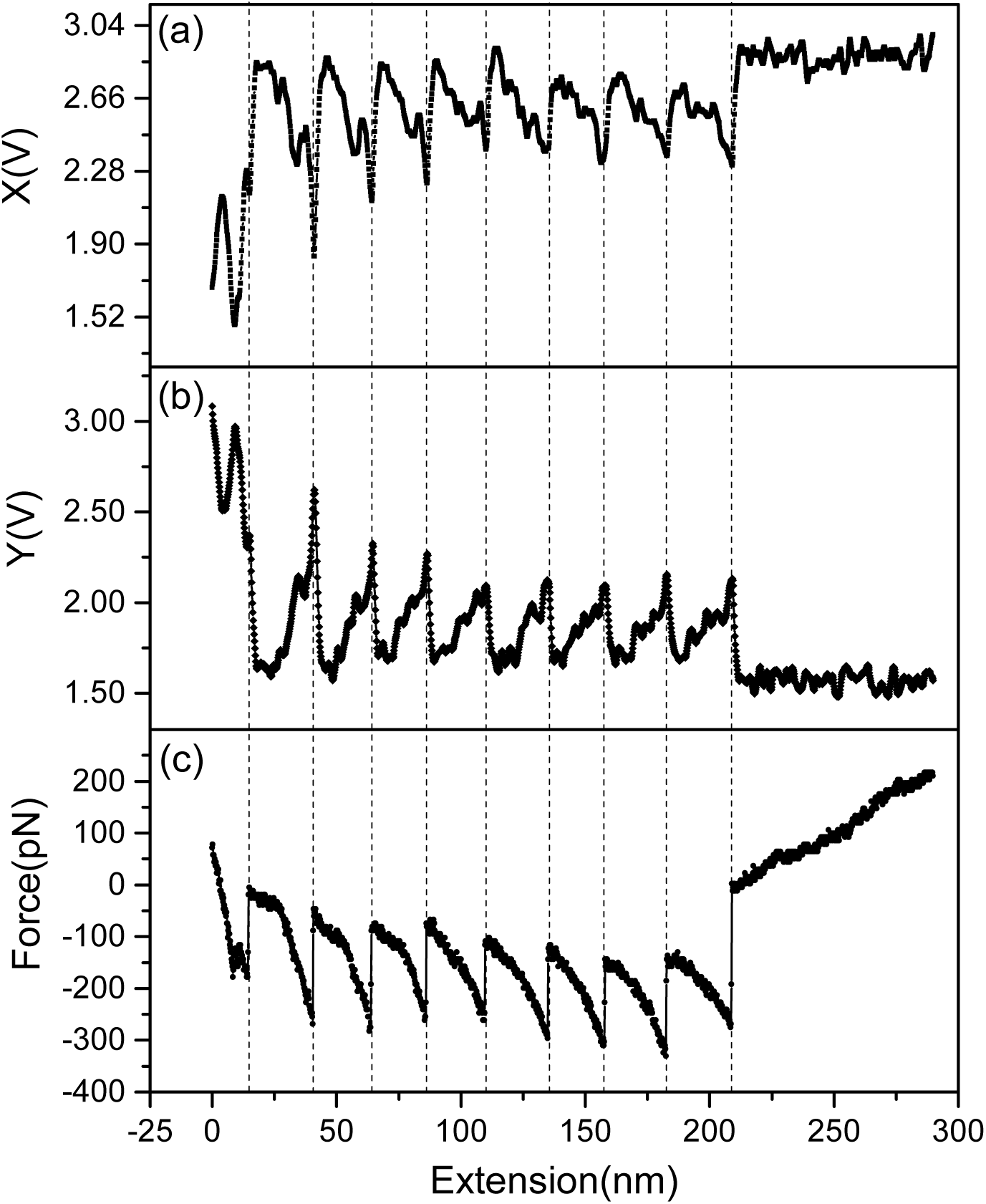
The oscillatory motion of the cantilever can be split into two components. One component is in phase and the other is 90 degree out of phase with the drive. These are termed the X and Y components of motion respectively. The resultant of these two components is the tip amplitude; *R*^2^ = *X*^2^ + *Y* ^2^. The phase angle between the drive and the cantilever end bearing the tip is *θ* = *tan*^−1^(*Y/X*). a) x-component (*X*) and b) y-component (*Y*) of the motion directly acquired from the lock-in amplifier. According to the beam theory used by Benedetti et al., the *X* and *Y* components exclusively contain the stiffness and dissipation information whereas the amplitude and phase data each contains both. c) Simultaneously acquired force-extension data, the conventional force curve in the literature.

### The stiffness and dissipation

To analyse stiffness and dissipation data, it should be noted that each domain of poly-protein can be considered as spring and dashpot in parallel. This means that dissipation from all domains add in a manner similar to springs in series. The lowest dissipation and stiffness among all domains is measured in experiments. Assuming that the stiffness and dissipation of unfolded domain is less compared to the folded domains, dissipation and stiffness measured here is that of an unfolded domain. See Supplementary information for details.

Figure 3 (a) shows stiffness calculated from data in figure 2 (a) and using equation 2. The stiffness is in the range of 10 −100 mN/m. It clearly shows a feature - a drop in stiffness before the peak stiffness is reached. This is marked by a star in figure 2 (a). To interpret the stiffness data in our measurement it is critical to understand the process of unfolding of domains of I27 upon application of force. In past, AFM measurements have shown that unfolding of I27 occurs in two steps which reveals an intermediate characterised by elongation of folded domains by around 15 percent. It is understood through SMD that this is a result of breaking of two H-bonds between A and B *β*-strands. Further pulling breaks six H-bonds between A’ and G which unfolds the protein completely. In static measurements, the signature of this intermediate is a small plateau in otherwise monotonous force-extension curves following a Worm-Like-Chain behaviour. In our experiments the measured stiffness is that of the weakest spring in the system. After a domain unfolds, the stiffness of its entropic chain is lowest compared to remaining folded domains. Upon further stretching, the entropic stiffness increases following a Worm-Like-Chain. This transmits the applied force effectively to folded domains which allows application of ~ 100 pN of force required to push the folded domains into an intermediate state with elongation along the direction of pulling. The drop in stiffness at ∼100 pN force in our measurements corresponds to this well known intermediate of I27. At this point the measured stiffness is that of an unfolded domain and remaining folded domains in their intermediate state. Further stretching increases stiffness of domains in intermediate and entropic part of the unfolded domain dominates the measurement and follows WLC. We fitted a derivative of Marko-Sigga approximation to stiffness-extension curve. The WLC fits on both peaks returned two contour lengths with same persistence length (0.38 ± 0.04 nm). Difference in contour lengths (Δ*L*) is the additional extension coming from elongation of remaining folded domains. Δ*L* increases with number of folded domains which indicates reversible folding of the intermediates upon removal of force. Figure 3 (b) shows a plot of Δ*L* versus the number of folded domains. The slope of this line is elongation per domain. The average elongation estimated from the slope is ∼ 8 Å per domain, consistent with previous reports and SMD [44].

**FIG. 3:**
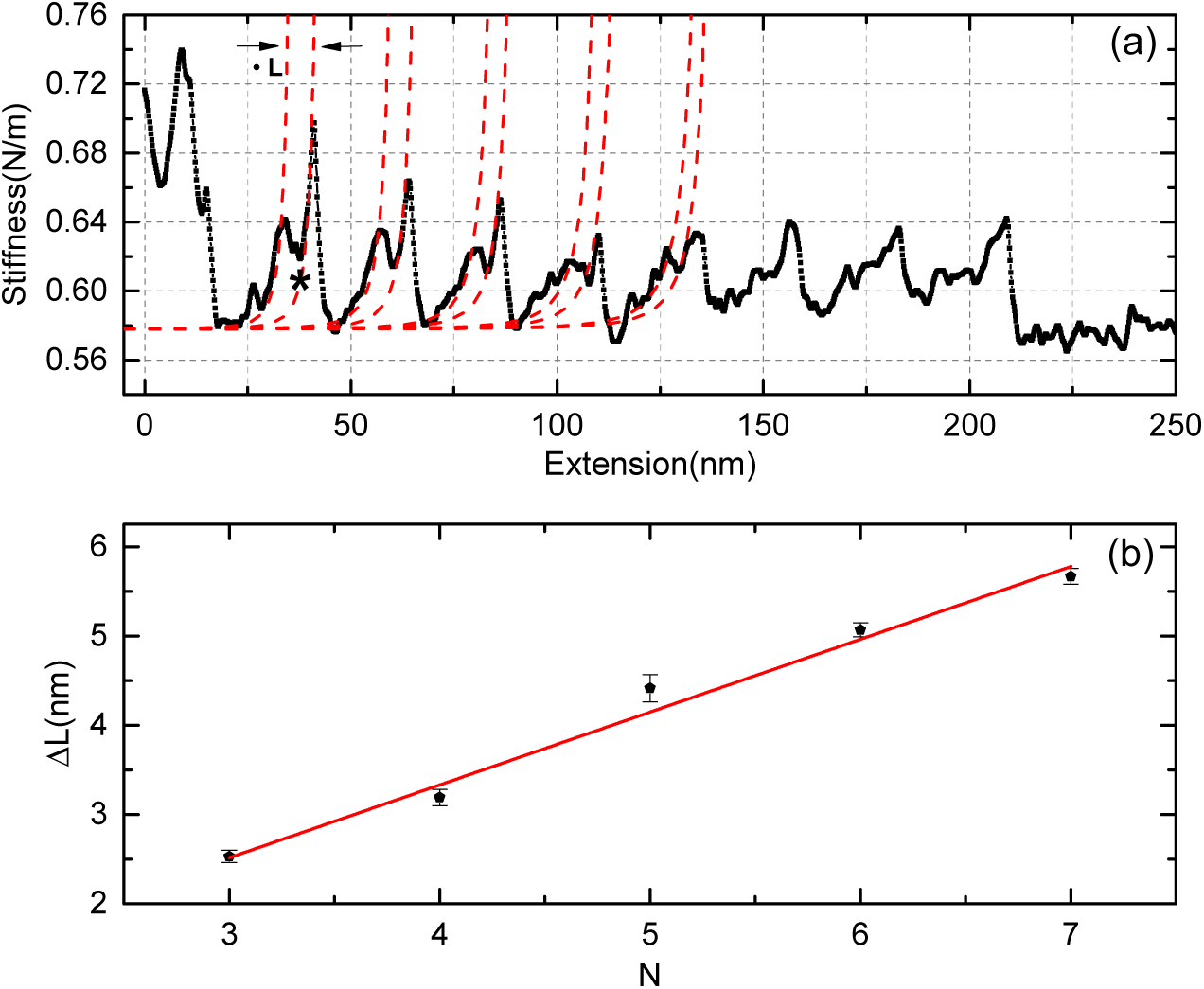
(a)Stiffness calculated from X data in figure 2(a) and using equation 2. The peaks in stiffness data correspond to peak force in force-distance curve of figure 2(c). The measured stiffness is that of the weakest spring in the series. It is the unfolded domain and has a mechanical response similar to a random polymer chain in coiled state. It follows strain-stiffening governed by a Worm-Like-Chain model (dotted lines). See the methods section. As the polymer chain stiffens, it causes the folded domains to fall into an intermediate which has similar or less stiffness than the entropic stiffness of the unfolded domain (marked by star). Increasing the strain further enhances the stiffness of the chain following WLC. We fitted WLC model to stiffness-extension data both before and after the remaining domains fall into an intermediate. The intermediates produce extra extension Δ*L* by elongating the domains. This can be obtained by subtracting the contour lengths of two fits with same persistence length (0.38± 0.04 nm). (b) Plot of Δ*L* versus no. of remaining folded domains N. The slope of the line is elongation per domain after the breakage of two hydrogen bonds between two of the *β* strands.

Figure 4 (a) shows dissipation computed using equation 3 on data in figure 2 (b). Although, we follow method by Benedetti et al. for computing dissipation from the Y-data, we see that the dissipation in our measurement is above the detection limits. The measured dissipation is attributed to the internal friction of unfolded domains. Through fluctuation measurements, it is shown that the dissipation in unfolded globule increases with tension following a power law. Such a globule is considered to be dry but devoid of native contacts and not having 3D conformation of a native folded state. The WLC model with consideration of internal friction due to dihedral rotations predicts a scaling with force ∼ *F* ^1/2^ [17]. The dissipation in unfolded domain increases with force and the measured dissipation drops after sudden drop in force at each unfolding event. Interestingly, and for the first time, our dissipation data shows an abrupt decrease corresponding to intermediate signal in stiffness data. Since force on the protein does not change, the drop in dissipation is not a result of reduced tension on the unfolded entropic polymer chain. It implies that the transition from native structure to folded intermediate has caused the internal friction in these domains to become less than the unfolded part and hence contribute to the measurement. The value at this point is combined dissipation of an unfolded chain and folded intermediates of I27 resulting from the breakage of two hydrogen bonds between A and B *β* strands. Further, we have chosen data which does not have the intermediate signature in the dissipation. As figure 4 (a) suggests, this occurs when there are less number of folded domains remaining of the stretched poly-protein. Figure 4 (b), a log-log plot of dissipation versus force, reveals that indeed the dissipation scales with force and the exponent is 1/2. It indicates that our measurements are largely affected by dihedral rotations alone.

**FIG. 4:**
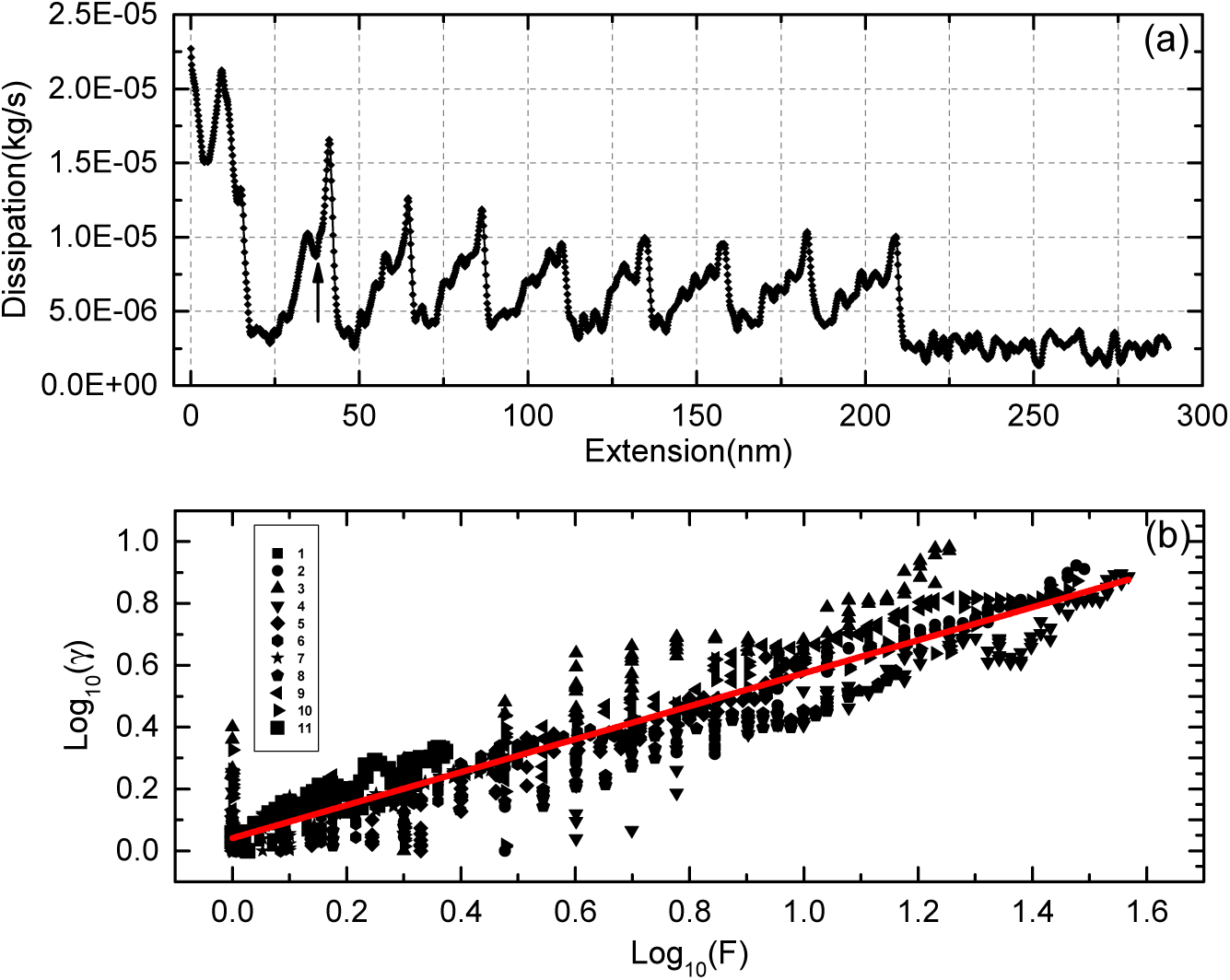
(a) Dissipation calculated from Y data in figure 2 and using equation 3. The peak in dissipation corresponds to the peak force in force-extension curves. When a domain unfolds the dissipation is lowest and increases as the unfolded domain is stretched further. When the dissipation from elongated intermediates is lower than that of the unfolded domain, a drop in dissipation is seen (marked by an arrow). At this value of tension in the unfolded domain, its dissipation is higher than intermediates. b) Dissipation versus force data from unfolding of individual domains. The symbols are experimental data and the red continuous line is fit with exponent 1/2.

## DISCUSSION

Internal friction plays a significant role in determining protein folding dynamics[27, 48]. In past, it was inferred indirectly from reconfiguration time-scales in protein folding using fluorescence-based and other methods[46]. Internal friction in single molecules can be probed directly by force measurements having pN to nN sensitivity. With advent of single molecule techniques, such as AFM and optical tweezers, it has been possible to measure internal friction. Recently, doubts have been raised over suitability of small amplitude AFM to detect internal friction in the unfolded protein domains[28]. We clearly show here that dissipation resulting from internal friction can be directly probed with AFM. While Benedetti et al. are correct in pointing out a gross oversimplification of cantilever dynamics in liquid environment, their conclusion, that the internal friction is below detection limit of AFM, is based on experiments carried out with unsuitable parameters (See supplementary information). Indeed, the approach of small amplitudes (∼ 1 Å), large cantilever stiffness (∼ 1 N/m) and purely off-resonance operation is essential for dissipation measurement using small amplitude AFM[35]. This has also been the case for probing the dissipation in layering of liquid molecules next to a solid wall using small amplitude AFM [41, 47].

Kawakami et al. used resonant AFM with large amplitudes (∼ 2 nm) to measure stiffness and dissipation of unfolded domains by approximating the cantilever with a point-mass. They note that there is no abrupt change in dissipation corresponding to unfolding intermediate feature in stiffness data. It led them to conclude that the internal friction of native state and the folded intermediate is similar. As noted in figure 3 and discussed in previous paragraph, we observe a sudden decrease corresponding to the intermediate. It means that while measuring viscoelastic response of single molecules using AFM, choice of analysis method as well as employment of amplitudes smaller than persistence length (≤ 3 Å) is important to clearly observe the dissipation features which are otherwise lost in poor S/N ratio.

How general are the values (∼ 10^−5^ kg/s) obtained for single molecule internal friction in our experiments? The friction coefficient measured using fluctuations of an unfolded chain at high stretch (*fb* > *K*_*B*_*T*; where f-Applied force, b-persistence length) have yielded values in the same order. Wang and Zocchi have measured friction coefficient of folded proteins using AC susceptibility[49]. It is 7.8 × 10^−5^ kg/s per molecule. Kawakami et al. conclude that friction of folded native structures and intermediate have a similar value. Together with our measurements, these observations point out that internal friction in folded, unfolded or partially folded domains is of the order of 10^−5^ kg/s. This suggests that role of native contacts in determining internal friction is not significant. Our observation, that the folded intermediate is less dissipative compared to stretched unfolded domain, suggests a need for better understanding of origins of internal friction in folded as well as unfolded proteins. Assuming length scale of *a* = 10 nm for single protein domain, a typical value of solvent friction with unfolded domain would be *γ*_*hydro*_ = 6*πηa* ∼ 10^−11^*kg/s*, many orders smaller than internal friction measured here.

Phase sensitive detection techniques such as lock-in amplifiers and phase lock-loops are often used for detecting signal which is weaker compared to noise. It is clear from our measurements that when static force signal is not able to show presence of intermediates(figure 2 c), the stiffness data obtained from use of lock-in amplifier shows its presence unambiguously (figure 2 a and figure 3 a). This was also demonstrated in frequency shift measurements using Phase-Lock-Loop by Higgins et al. [50]. We show that an additional advantage of small amplitude approach over resonant frequency tracking is its ability to measure friction coefficient of unfolded domains. In many proteins, the chemically induced unfolding and static AFM data has failed to decipher intermediates and shows two-state, all-or-none transition, but force clamp methods on the same proteins have shown unfolding through intermediates [51–53]. Direct measurement of stiffness using small amplitude AFM should be a method of choice for detecting the presence of intermediates along the unfolding pathway for these proteins.

Benedetti et al. have criticised use point-mass model to determine dissipation and stiffness using phase and amplitude. The point raised by them about appropriate cantilever modelling is important and it is previously addressed by many others[38, 39]. However, there are reports of dissipation measurement of single molecules using AFM, wherein the cantilever is driven thermally and phase is not measured to estimate the dissipation[17]. Such measurements have revealed that the internal friction of single molecules, both proteins and polysaccharides, is of the order of ∼ 10^−5^ kg/s. It is larger than the hydrodynamic dissipation coefficient of the cantilever itself (∼ 10^−6^ kg/s), which has many orders of magnitude (∼ 10_4_) more area exposed to the water compared to the protein.

## CONCLUSIONS

In conclusion, we measured dissipation resulting from internal friction of single unfolded domains in I27 using small amplitude AFM. The stiffness data clearly shows presence of intermediates in titin domains as predicted by molecular dynamics simulations in the past. We demonstrate that small amplitude AFM approach is more suitable for measuring viscoelastic response of single molecules. The internal friction in single molecules competes with hydrodynamic dissipation coefficient of cantilever itself in liquids and is order of magnitudes larger than the solvent friction with unfolded protein domain. The measurements clearly underline the importance of appropriate operational parameters for detecting dissipation in single molecules using small amplitude AFM technique. It can be a useful tool to directly probe differences in internal friction of unfolded *β* sheet rich (titin) and *α* helix rich (*α*-spectrin) protein domains. In future, such measurements may shed light on the origins of folding rates of different tertiary structures.

## AUTHOR CONTRIBUTIONS

ST, SR and VA collected data, performed analysis and prepared figures. SK and AR made protein samples. VA and SR helped in writing the manuscript. SP conceived experimental protocol, analysed data and wrote the manuscript.

## ACKNOWLEDGEMENT

The work is supported by Wellcome Trust-DBT India Alliance (500172/Z/09/Z). The authors would like to thank ASR Koti from Tata Institute of Fundamental Research for providing the plasmid. VJ and VA would like to thank DST INSPIRE for fellowship. SP acknowledges the Intermediate Fellowship awarded by Wellcome Trust-DBT India Alliance.

## Supplemental Materials: Direct Measurement of Dissipation in a Single Protein using Small Amplitude Atomic Force Microscopy

### Spring and dash-pot representation of the polyprotien and cantilever

**FIG. S1:**
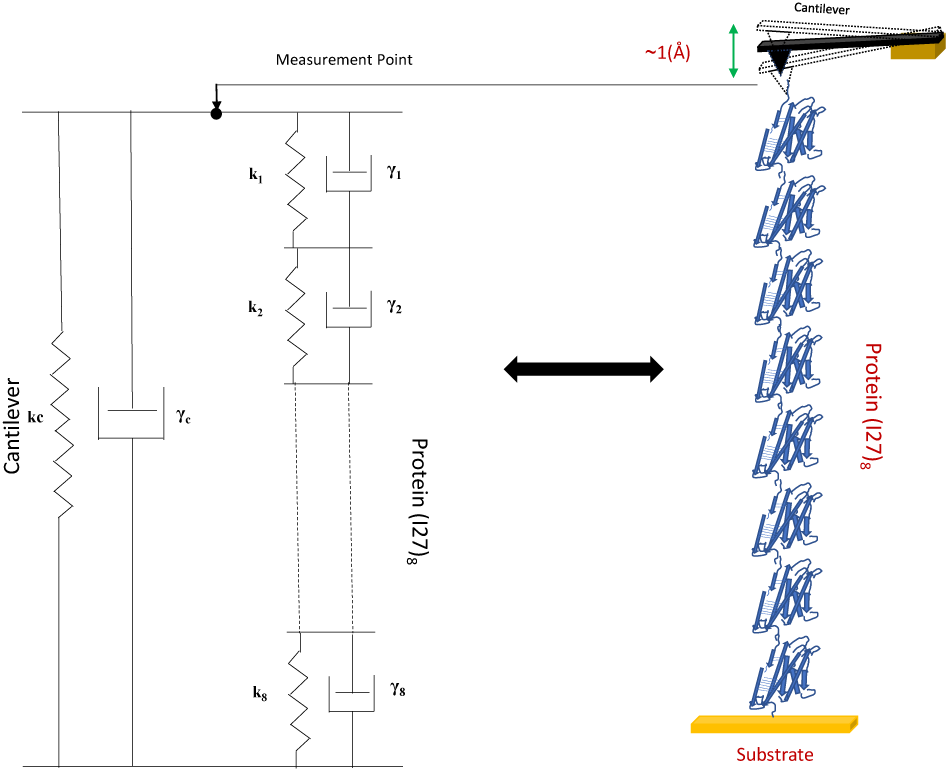
The spring and dash-pot model representing protein polymer stretched using an AFM tip. The 8 monomers are arranged end to end to form the polymer. This can be viewed as 8 springs and dash-pots in series. The ends of each dash-pot and springs are connected together. The spring representing the cantilever is parallel with protein spring. Similarly the dash-pot representing the hydrodynamic drag on the cantilever is in parallel with protein dash-pots.

Fig.S1 shows a schematic of the measurement of dissipation and stiffness in a single polyprotein consisting of 8 monomers using small amplitude AFM. The protein is tethered to end of the cantilever and gold substrate. The cantilever is oscillated with small amplitudes (∼ 1 Å) and is steadily pulled away from the substrate with a constant velocity(∼ 50 nm/s). The polyprotein is represented as eight springs and dashpots in series. The springs and dashpots representing individual domains are in parallel, since the displacement at these points is same. The spring and dashpot representing the cantilever stiffness and damping coefficient is parallel to polyprotein, since the measurement is being performed at the tip. The resulting equations for stiffness and dissipation are given below,

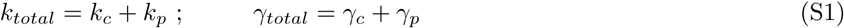

Where *k*_*c*_ and *γ*_*c*_ are cantilever stiffness and dissipation coefficient respectively. The cantilever dissipation is due to the hydrodynamic damping provided by the surrounding medium. The springs and dashpots representing individual domains add together in following manner.

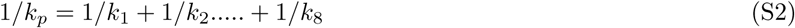

and

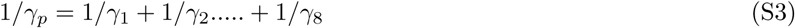

Where *k*_1_ …. *k*_8_ etc. are stiffness and *γ*_1_ …. *γ*_8_ etc. are damping coefficients of individual domains. For identically prepared domains such as octamer of I27, stiffness and dissipation of folded domains are equal to each other; *k*_1_ = *k*_2_ = *…….k*_8_. Similarly, for damping coefficient *γ*_1_ = *γ*_2_…………*γ*_8_. In small amplitude AFM measurements when one measures *k*_*p*_ or *γ*_*p*_, the measurement is dominated by their lowest values. Typically, after a given domain unfolds, its stiffness drops considerably for low strain values and increases with high strain following WLC model. Similarly, lowest dissipation of dash-pots in series is recorded in experiments. The drop in stiffness and dissipation when folded domains fall into intermediates suggests that at high stretch, the dissipation of intermediates is less than the unfolded polymer chain.

### Measurement of dissipation in single molecules with small amplitude AFM

Benedetti et al., using small amplitude AFM, claimed that it is not possible to measure dissipation in single molecules with this technique. They concluded that the dissipation in single molecules both, internal and due to hydrodynamic coupling, is smaller than the AFM detection limit. This was estimated to be about 10^−7^*kg/s*. The dissipation measured in present work is ∼10^−5^ kg/s. Typical thermal noise floor of cantilevers used in AFM measurements are same. The detection sensitivity of commercial AFM systems is of the same order of magnitude. What is the reason behind this apparent discrepancy ?

We could not do measurements at exactly the same parameters as used by Benedetti et al. for technical reasons. However, in the following we show that choice of operational parameters such as lever stiffness, resonance frequency, operational frequency and amplitude play a crucial role in successfully measuring dissipation. First, we show that stiff cantilevers used in the current measurement may not show variation in Y (dissipation signal) even as protein unfolds sequentially. If the operation frequency is large (more than *ω*0*/*3), the dissipation signal related to single molecules unfolding is not observed. Figure S2 shows measurements performed with a cantilever having similar stiffness which is used record data in figure 2 in the main text, but we do not drive the cantilever in strictly off-resonance condition. The resonance frequency of the lever is 6.7 KHz. Instead of using drive frequency which is far below resonance, we used a larger frequency ∼3.5 kHz. Now the cantilever dynamics is complicated and it is not fully captured in theoretical treatment which assumes off-resonance condition. The Y component in figure S2 does not show any feature. We performed measurements with weak cantilevers (0.02 N/m). This is the typical cantilever stiffness used to measure the unfolding force in static experiments and also used by Benedetti et al. We found that for small amplitudes (∼ 1Å) and low frequencies (140 Hz), we could observe the Y signal in lock-in amplifier, but it was noisy. The experiments at higher frequency (900 Hz) and larger amplitudes (∼8Å) were again featureless.

The mathematical treatment used to calculate dissipation and stiffness using small amplitudes relies strongly on the off-resonance approximation. If the experiments are not conducted off-resonance the Y signal may not represent the dissipation and one may incorrectly interpret this as dissipation is below the limits of detection. As this is shown amply in all previous small amplitude works, it is quite important to work with stiff cantilevers, small amplitudes and off-resonance to use the method correctly [S35]

**FIG. S2:**
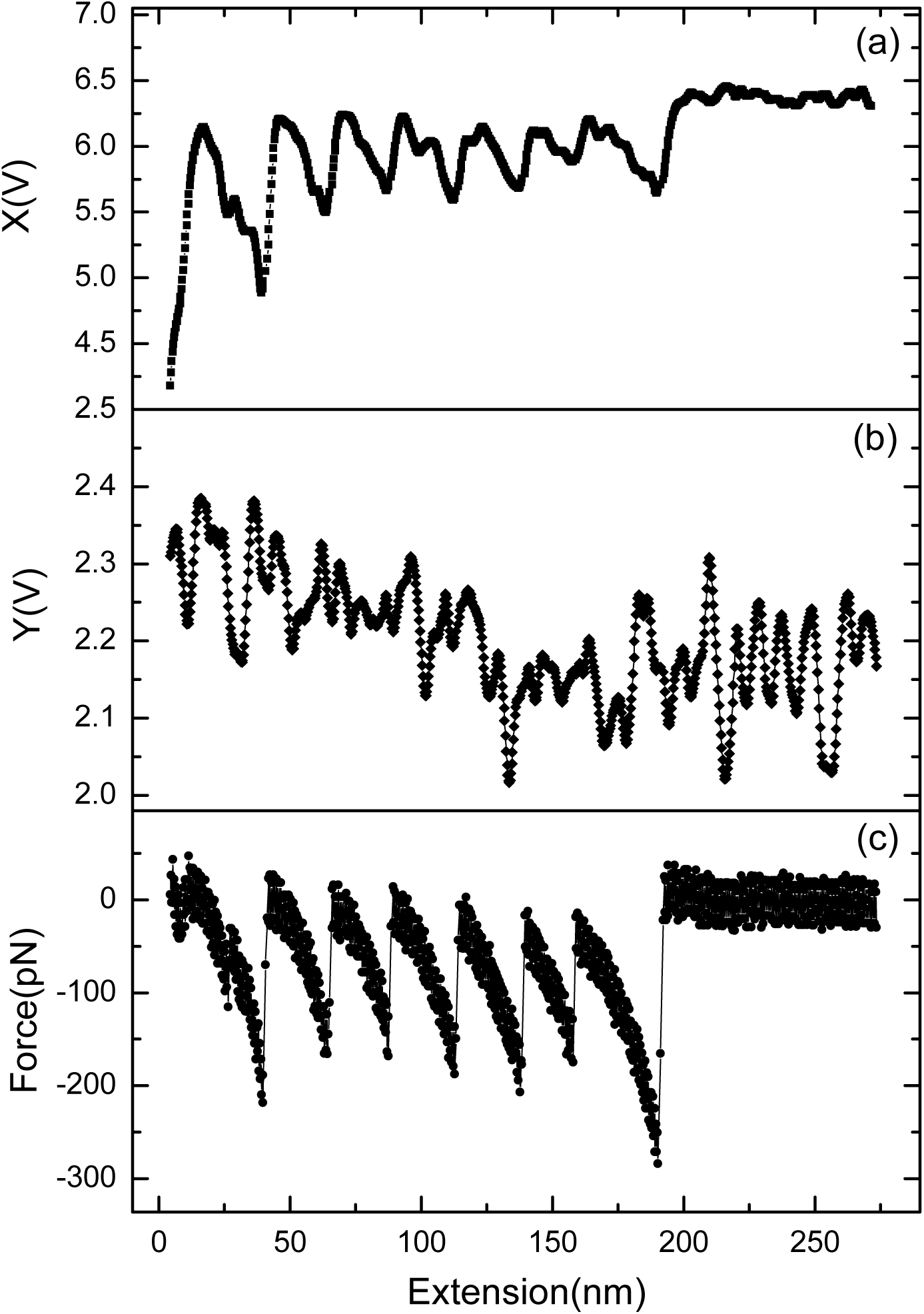
Raw data of pulling experiments performed with stiff levers (0.2 N/m) but not strictly off-resonance where drive frequency is only half of the resonance frequency (*ω*_*d*_ ∼ *ω*_0_*/*2. The amplitude is 8.4 Å. The Y-signal, which corresponds to dissipation does not have features as protein domains undergo transition from folded to unfolded state.

**FIG. S3:**
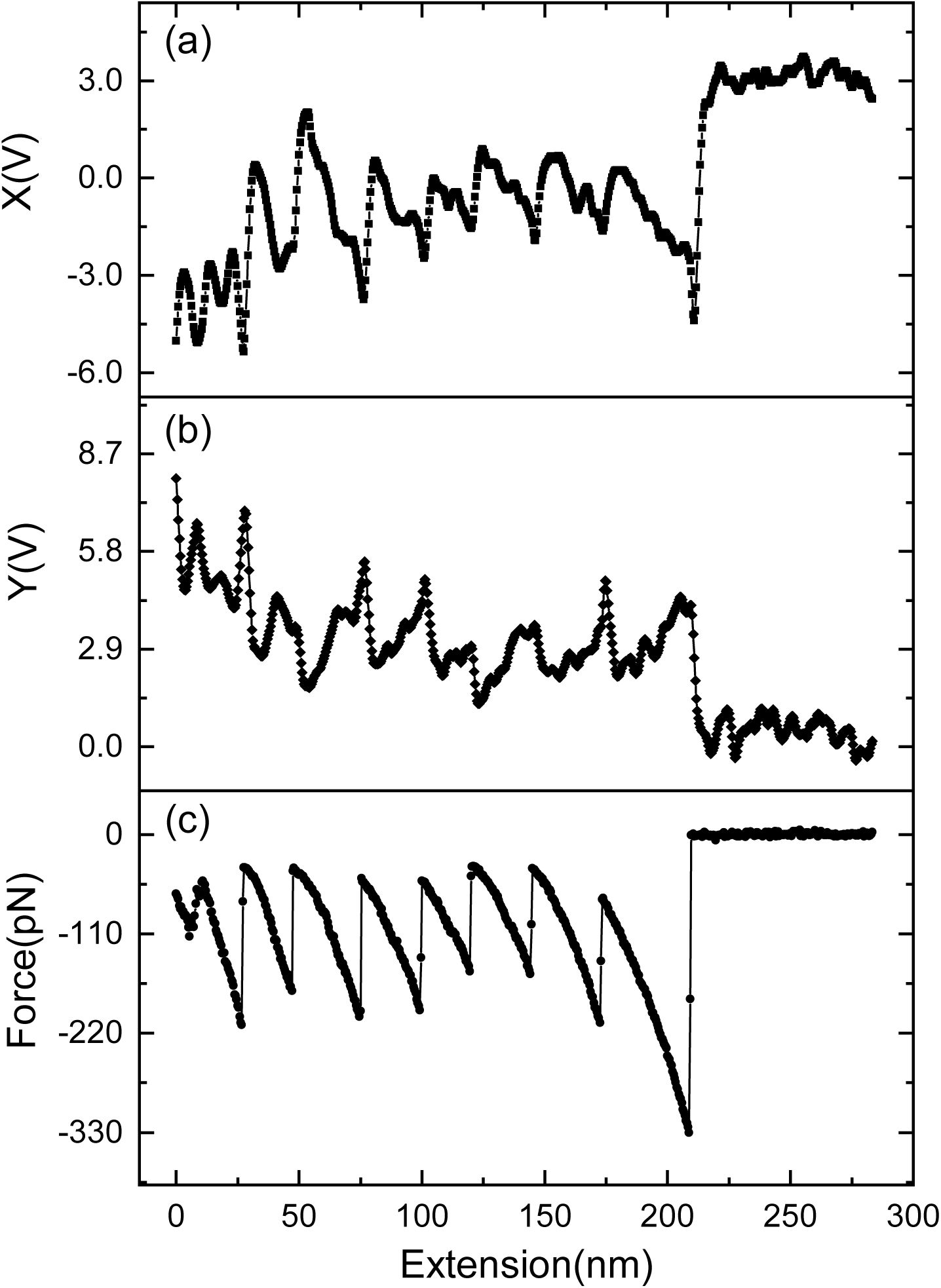
Raw data of pulling experiments with small amplitudes (∼ 1.3 Å) and weak levers (0.02 N/m) and off-resonance operation (*ω*_*d*_ = 140 Hz, *ω*_0_ = 1.5 KHz). There are features with certain noise in the Y-signal. This indicates that weak levers, small amplitudes is able to measure dissipation signal, However thermal noise in case of weak levers produces low signal-to-noise ratio.

**FIG. S4:**
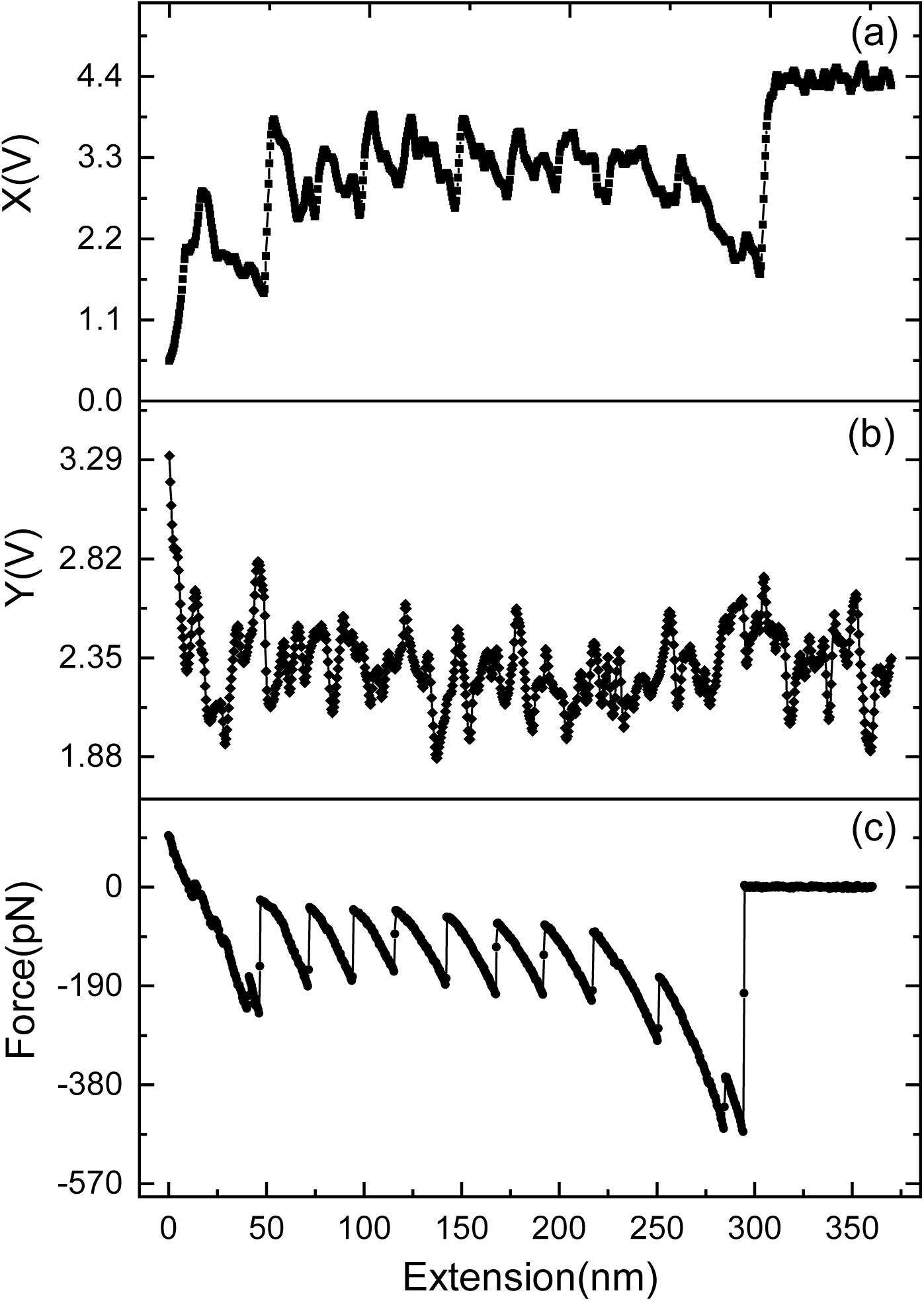
Raw data of pulling experiments with weak cantilevers (0.02 N/m) and large amplitudes (∼ 8Å) but strictly not off resonance. *ω*_*d*_ = 900*Hz* and *ω*_0_ = 1.5 KHz. The Y signal does not show features related to protein unfolding.

